# ATP triggers macropinocytosis that internalizes and is regulated by PANX1

**DOI:** 10.1101/2020.11.19.389072

**Authors:** Andrew K.J. Boyce, Emma van der Slagt, Juan C. Sanchez-Arias, Leigh Anne Swayne

**Affiliations:** Current affiliation: Hotchkiss Brain Institute; Department of Cell Biology & Anatomy; University of Calgary; Calgary, Alberta T2N 4N1, Canada; Division of Medical Sciences; University of Victoria; Victoria, British Columbia V8P 5C2, Canada

## Abstract

Macropinocytosis is an endocytic process that allows cells to respond to changes in their environment by internalizing nutrients and cell surface proteins, as well as modulating cell size. Here, we identify that adenosine triphosphate (ATP) triggers macropinocytosis in murine neuroblastoma cells, thereby internalizing the ATP release channel pannexin 1 (PANX1) while concurrently increasing cross-sectional cellular area. Amiloride, a potent inhibitor of macropinocytosis-associated GTPases, abolished ATP-induced PANX1 internalization and cell area expansion. Transient expression of the GTP-hydrolysis resistant GTPase ARF6 Q67L led to increased PANX1 internalization and increased cell area equivalent to levels seen with ATP stimulation. Mutation of an extracellular tryptophan (W74) in PANX1 abolished ATP-evoked cell area enlargement suggesting that PANX1 regulates this form of macropinocytosis. This novel role of PANX1 in macropinocytosis could be particularly important for disease states implicating PANX1, such as cancer, where ATP can act as a purinergic regulator of cell growth/metastasis and as a supplementary energy source following internalization.

## INTRODUCTION

Amongst the many ways that cells adapt to environmental changes, several key modifications include cell surface receptor density, nutrient acquisition, and cell size. Macropinocytosis, or “cell drinking,” is a non-canonical membrane internalization process that enables these tightly-associated dynamic adaptations [1]. In some cell types, macropinocytosis enables recycling of the equivalent of the full cell surface area once every thirty minutes [2]. A more thorough appreciation of the regulation of macropinocytosis is important for understanding antigen presentation in phagocytic immune cells [3] and receptor-mediated signalling localized to endosomal compartments [4,5], as well as many types of virus and pathogenic bacteria entry [6–9] and nutrient acquisition in tumorigenic cells [10,11] (for recent reviews see Finicle et al. [12] and Bloomfield and Kay [13]).

In this report, we investigate the hypothesis that ATP-mediated PANX1 internalization, a phenomenon we recently discovered [14,15], occurs through macropinocytosis. Macropinocytosis is distinct from classical types of membrane internalization or endocytosis in that it does not rely on coat and scission proteins used in clathrin- and caveolin-mediated endocytosis. Instead, during macropinocytosis, discrete regions of cholesterol and phosphatidylinositol-4,5-bisphosphate (PI(4,5)P_2_)-rich plasma membrane and its associated receptors are engulfed via actin-dense membrane ruffles [16]. Initially, ADP-ribosylation factor 6 (ARF6), a small GTPase resident to the membrane ruffles, activates phosphatidyl inositol-4-phosphate 5-kinase [17] to generate PI(4,5)P_2_. Generation of PI(4,5)P_2_ enables recruitment of the small GTPases Ras-related C3 botulinum toxin substrate 1 and cell division control protein 42 (more commonly known as CDC42) to the ruffling membrane [18,19], which then activate p21-activated kinase [20]. Next, p21-activated kinase phosphorylates a protein called brefeldin A-dependent ADP ribosylation substrate to initiate membrane curvature [21], which is enabled by conversion of lysophosphatidic acid to phosphatidic acid (for review see Bohdanowicz and Grinstein [22]), leading to the formation of large vesicles ranging in diameter from 0.2 to 5 μm referred to as macropinosomes. Given the lack of coat protein, structural specificity, or unique membrane-bound molecules, macropinosomes are difficult to distinguish from other endocytic compartments of similar size, and consequently, are commonly identified using fluorescently labelled fluid-phase endocytosis markers (*i.e.*, FITC-dextrans) [23]. Fortunately, the small GTPases that regulate macropinocytosis are uniquely sensitive to changes in sub-membranous pH; and for this reason, amiloride inhibition of the Na^+^/H^+^ exchanger selectively targets macropinocytosis over other forms of endocytosis [24].

Depending on the cell type, macropinocytosis can be a constitutive process, or can be triggered by extracellular stimuli, [25]. Examples of cells that undertake constitutive macropinocytosis include immature dendritic cells (for sampling of soluble antigens [26,27]) and cancerous fibroblasts transformed with oncogenic K-Ras or v-Src [28]. Extracellular molecules that can stimulate macropinocytosis include growth factors (e.g. epidermal growth factor [29], macrophage colony-stimulating factor [30]), and chemokines [31]; this breadth of stimuli enables context-dependent regulation of signalling at the cell surface. Although not a canonical growth factor, adenosine triphosphate (ATP) plays a growth factor-like role in regulating many cellular functions through activation of purinergic receptors [32–35]. ATP is released via a number of mechanisms including through channels like pannexin 1 (PANX1) as well as by vesicular release (reviewed in [36,37]). PANX1 forms ubiquitously-expressed heptameric channels that are permeable to small (Cl^−^) and large (e.g. ATP) anions [38–44]. Additionally, PANX1 may also regulate the flux of small cations (Ca^2+^) [45] and release of large cations (spermidine) [46].

Selectivity for anions or cations has been proposed to depend on the mechanism of channel activation [47]. PANX1 anion selectivity, characteristic of certain open states [38,43,44], has been attributed to extracellular tryptophan (W74) and arginine (R75) residues that form a molecular filter for size and charge, respectively. While their overall properties are still the focus of intense investigation, it has been well-established that PANX1 channels play a role in ATP release [36,48–54]. Once in the extracellular space, ATP, as well as its metabolites arising from the activity of tissue-specific ectonucleotidases [55] trigger an array of downstream signaling pathways that can lead to myriad cellular changes.

We and others have demonstrated that extracellular ATP can also directly impact the ATP release machinery, thus effecting sustained changes in cell signalling [14,56–58]. Exogenously added extracellular ATP can inhibit PANX1 channel activity [57–59] as well as stimulate channel internalization [14,60]. Cell surface physical association of PANX1 with the ionotropic purinergic P2X7 receptor (P2X7R) preceded internalization [14]. Site-directed alanine substitution of the PANX1 extracellular loop tryptophan, W74, disrupted cell surface P2X7R - PANX1 association, as well as PANX1 internalization [60], suggesting this residue regulates both ionic selectivity and cell surface stability. In the course of this work, we noted that internalized PANX1 was localized to structures much larger than those characteristic of canonical endocytic pathways [14]. ATP-triggered internalization of PANX1 did not utilize the canonical clathrin-, caveolin-, and dynamin-associated machinery, consistent with work from other groups [61,62]. Furthermore, ATP-evoked PANX1 internalization in N2a cells coincided with active filopodial dynamics, relied on cholesterol [14,60], and occurred in the absence of canonical endocytic effectors, suggesting a non-canonical mechanism, such as macropinocytosis.

We first examined the impact of the macropinocytosis blocker amiloride [24] on ATP-induced PANX1 internalization. As anticipated, amiloride, prevented ATP-evoked PANX1 internalization. Additionally, expression of a constitutively-active ARF6 GTPase [17,18] increased basal intracellular PANX1 levels and prevented further ATP-induced PANX1 internalization suggesting that the macropinocytosis effector plays a role in ATP-induced PANX1 internalization. Finally, PANX1 interacted with specific phospholipids important for recruiting macropinocytosis effector proteins and generating membrane curvature during macropinocytosis, including PA, phosphatidylinositol-4-phosphate PI(4)P, and PI(4,5)P_2_ [21,63]. Consistent with expected cell volume changes following macropinocytosis [64], ATP induced a concurrent increase in cross-sectional cell area. This cell area expansion was disrupted by amiloride, and expression of PANX1 W74A. Taken together, these results suggest that, in addition to undergoing macropinocytosis in response to extracellular ATP, PANX1 could also play a broader role in the regulation of ATP-regulated macropinocytosis.

## METHODS

### Plasmids

The PANX1-EGFP and PANX1-RFP plasmids[65] were generous gifts from Drs. Dale Laird and Silvia Penuela. The pARF6-CFP (Plasmid #11382), pARF6 Q67L-CFP (Plasmid #11387), and pARF6 T27N-CFP (Plasmid #11386) were acquired from Addgene, courtesy of Dr. Joel Swanson [65].

### Cell Culture

Neuro2a (N2a) mouse neuroblastoma cells (procured from the American Type Culture Collection, ATCC, in 2011) were cultured in Dulbecco’s modified Eagle’s medium (DMEM)/F12 supplemented with 10% FBS, 100 units/mL penicillin, and 100 mg/mL streptomycin (all obtained from Gibco/Life Technologies). Where indicated, N2a cells were transfected using jetPEI reagent (Polyplus transfection/VWR) according to the manufacturer’s protocol. N2a cells stably expressing PANX1-EGFP or PANX1-RFP were maintained in DMEM/F12 containing 10% FBS, and 100 units/mL penicillin, 100 μg/mL streptomycin, and 400 μg/mL geneticin 418 (all obtained from Gibco/Life Technologies). Stable cell lines were generated as follows: N2a cells were transfected with PANX1-EGFP using the jetPEI reagent (Polyplus transfection/VWR) according to the manufacturer’s protocol. Cells were plated on poly-D-lysine (PDL, Sigma)-coated coverslips. For all internalization-related experiments, protein translation was briefly inhibited by treatment with 20 mg/mL cycloheximide (CHX; Sigma) for 8 h, coincident with other treatments. Cells were fixed with 4% paraformaldehyde (PFA) washed three time in phosphate buffered saline (PBS) prior to mounting for imaging or processing for immunostaining. We investigated the role of ATP (500 μM, Sigma) versus vehicle control (equal volume of water) on PANX1 cell surface stability and trafficking. We disrupted macropinocytosis by pre-treating cells with amiloride (300 μM; Sigma) or vehicle (DMSO) for 1 h.

### Immunocytochemistry

Antibody labelling of PFA-fixed cultures was performed as previously described [14,60]. The primary antibody used was early endosome antigen 1(EEA1 – 1:200; Cell Signaling) and the secondary antibody was Alexa647 AffiniPure donkey anti-rabbit IgG (1:600; Jackson ImmunoResearch).

### Imaging

Confocal imaging and analysis were performed blinded to the experimental conditions. Images were acquired with a Leica TCS SP8 confocal microscope. Quantification was performed using Leica Application Suite (version 3.1.3) and in the confocal plane displaying the largest plane of the nucleus (Hoechst 33342), where applicable. In the absence of Hoechst staining (where CFP-tagged constructs were expressed), a z-stack was captured over the entirety of the cells in the field of view and the middle z-plane was selected for ROI analysis. ROIs with a cross-sectional area of ≤ 60 μm^2^ were excluded from analysis (3 ROI total, 2 from ARF6-CFP vehicle, 1 from ARF6 Q67L-CFP – vehicle). Comparisons were made between images acquired under identical conditions. Representative confocal micrographs were adjusted for contrast uniformly using Adobe Photoshop (CC 2015.1.2) for display purposes only; no contrast adjustments were made prior to analysis. Confocal images (Leica TCS SP8) of fixed cells were acquired using a 40X (1.3 NA) oil immersion objective at 3X optical zoom in 1296 x 1296 format with a pixel area of 71 nm^2^ as confocal z-stacks. Quantification of PANX1-EGFP/PANX1-RFP fluorescence intensity to describe ‘intracellular PANX1’ was performed at time zero and at 30 min post stimulation, as follows: a polygonal trace was drawn 1 μm inside of the peak WGA intensity at the cell periphery and the encapsulated average PANX1-EGFP fluorescence intensity per pixel was computed (Figure 1). Quantification of cross-sectional area was determined on same z-plane as intracellular area. Cross-sectional area per cell was the area encapsulated by the region of interest (ROI) traced along the cell periphery, as described above.

**Figure 1.**
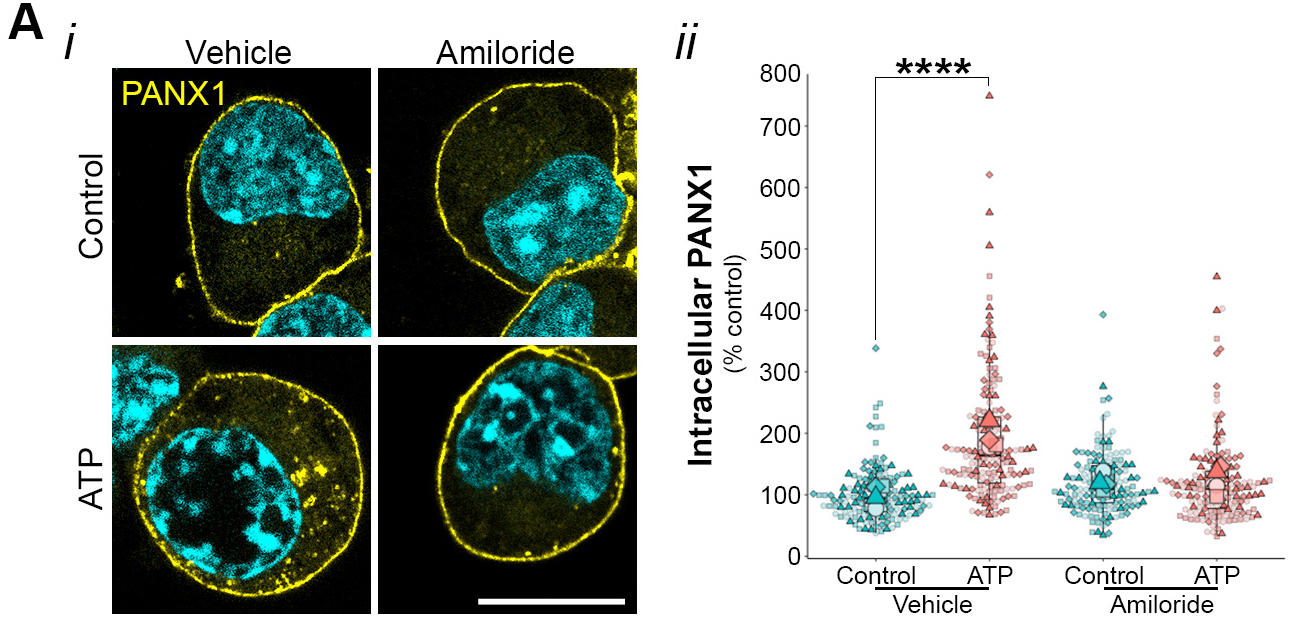
Amiloride disrupts ATP-induced PANX1 internalization. (**a**) Representative confocal micrographs of PANX1-EGFP (green) expressed in N2a cells (*i*) pre-treated with vehicle (equal volume of DMSO) or amiloride (300 μM) for 1 h prior to vehicle (water) or ATP (500 μM; 30 min). Hoechst (blue) was used as a nuclear counterstain. Scale bars, 10 μm. (*ii*) Quantification of intracellular PANX1 (normalized to control values). *N* = 4 coverslips from independent cultures (large symbols), technical replicates per coverslip are displayed in distinct shades of the same colour as smaller symbols. Data were subjected to one-way ANOVA with Dunnett’s *post hoc* (****p<0.0001). This figure was modified from the PhD thesis of A.K.J.B [67].

### Membrane Lipid Strip Interaction Assays

Membrane Lipid Strips (Echelon Biosciences; hydrophobic membrane spotted with 15 different membrane lipids [GT glyceryl tripalmitate, DAG – diacylglycerol, PA – phosphatidic acid, PS – phosphatidylserine, PE – phosphatidylethanolamine, PC – phosphatidylcholine, PG – phosphatidylglycerol, CL cardiolipin, PI – phosphatidylinositol, PI(4)P – phosphatidylinositol 4-phosphate, PI(4,5)P_2_ – phosphatidylinositol 4,5-bisphosphate, PI(3,4,5)P_3_ – phosphatidylinositol 3,4,5-triphosphate, CHOL – cholesterol, SM – sphingomyelin, SULF – 3-sulfogalactosylceramide] and a blank [xylene cyanol FF]) were blocked for 1 h in blocking buffer (3% BSA in PBS-T). Strips were then transferred to blocking buffer containing purified protein of interest: PI(4,5)P_2_ Grip (1 μg/mL, positive control from Echelon Biosciences), GST (5 μg/mL), or GST-fused PANX1 C-terminus (GST-PANX1^CT^, 5 μg/mL [66]) for 1 h, then washed three times in PBS-T and incubated for an additional hour in blocking buffer with primary antibody. Strips were again washed in PBS-T three times then incubated in corresponding HRP-conjugated secondary for 1 h. All incubations were performed at room temperature with gentle agitation.

### Study design and statistical analysis

The data presented are included in the PhD thesis of A.K.J.B [67]. The datasets used and/or analysed during the current study are available from the corresponding author on reasonable request. Data were normalized to values from vehicle treated PANX1-EGFP expressing controls. A biological replicate was defined as a coverslip obtained from an independent cell passage, where a technical replicate is each single cell on a given coverslip. Results were analysed using a one-way ANOVA or a Student’s t test, where applicable. Plots presenting superimposed summary statistics from repeated experiments (“SuperPlots” [68]) were generated using ggplot2, dplyr, and ggbeeswarm packages in RStudio (Version 1.3.1073). Data are presented as mean ± standard deviation (or mean ± S.E.M.; membrane lipid strips only). Detailed information about statistical tests is available in the figure legends.

## RESULTS

### Macropinocytosis inhibitor amiloride disrupted ATP-dependent PANX1 internalization

Macropinocytosis involves the coordinated recruitment and activation of several pH-sensitive GTPases in membrane ruffles [18,19]. Amiloride inhibits macropinocytosis by blocking the Na^+^/H^+^ exchanger resulting in acidification of the submembranous space and inhibition of requisite small GTPases [24]. We first tested the impact of pre-treatment with amiloride (1 hr; 300 μM) or vehicle (DMSO) on ATP-induced internalization of PANX1-EGFP constitutively expressed in murine neuroblastoma N2a cells (in the presence of transient protein-synthesis inhibition to reveal steady state changes in PANX1 localization) using an analysis paradigm established in our previous papers [14,60]. In the presence of amiloride pre-treatment, there was no change in intracellular PANX1 30 min post-stimulation with 500 μM ATP (Figure 1). This suggested that amiloride-sensitive macropinocytic GTPases were required for PANX1 internalization.

### Macropinocytosis effector ARF6 regulated ATP-dependent PANX1 internalization

ARF6 regulates the transit of cargo between the plasma membrane and a clathrin-independent endosomal compartment that can mature into a macropinosome[69,70]. We therefore assessed the role of ARF6 in ATP-dependent PANX1 internalization by co-expressing cyan fluorescent protein (CFP)-tagged wildtype ARF6, GTP-hydrolysis resistant (i.e., constitutively active) ARF6 Q67L, or ARF6 T27N (incapable of interacting with GTP), along with PANX1-RFP in N2a cells. As expected, ARF6Q67L expression triggered the formation and retention of large intracellular PANX1-positive/EEA1-positive vesicles independent of ATP stimulation (Figure 2A). Although dominant-negative ARF6 T27N is known to disrupt other forms of clathrin-independent endocytosis [71–74], there is little evidence for an effect on macropinocytosis; consistent with this, ATP-induced PANX1 internalization was not affected by co-expression of the wildtype ARF6 or the ARF6 T27N mutant (Figure 2B).

**Figure 2.**
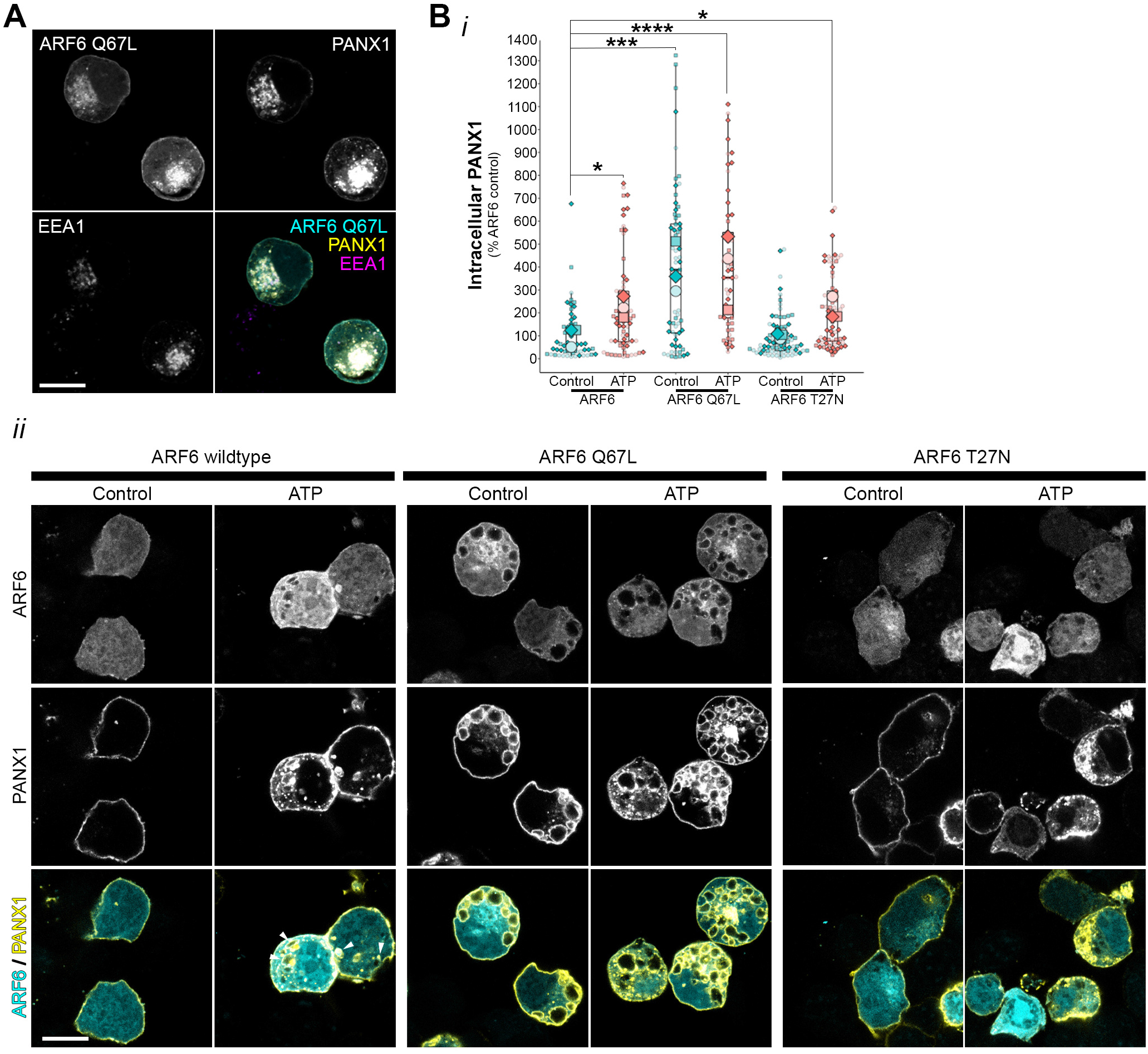
PANX1 is internalized in an ARF6-dependent mechanism. (a) Representative confocal micrographs of internalized PANX1-RFP (red) codistributed with ARF6 Q67L-CFP (green) and EEA1 (magenta) in intracellular structures of N2a cells. White arrows indicate overlap of internalized vesicles. (b) Quantification of intracellular PANX1 (red) following co-expression with wildtype ARF6-CFP, ARF6 Q67L-CFP, ARF6 T27N-CFP (green) following 30 minutes of vehicle (equal volume of water) or ATP (500 μM) treatment with representative confocal images of each condition presented in (c). Hoechst (blue) was used as a nuclear counterstain. Scale bars, 10 μm. *N* = 3 coverslips from independent cultures (large symbols), technical replicates per coverslip are displayed in distinct shades of the same colour as smaller symbols, data were subjected to one-way ANOVA with Dunnett’s *post hoc* (*p<0.05, ***p<0.001, ****p<0.0001). This figure was modified from the PhD thesis of A.K.J.B [67].

### PANX1 C-terminus interacts with lipids involved in macropinocytosis

The carboxy-tail of PANX1 (PANX1^CT^) contains a highly-disordered putative membrane-associated region [75] that was unresolved in recent cryo-EM structures (Figure 3A). As PANX1 internalization was cholesterol-dependent [14,60], we hypothesized that PANX1^CT^ interacts with cholesterol or other lipids commonly co-distributed with cholesterol or known to be involved in cholesterol-dependent processes. To test this prediction, we incubated purified GST-tagged PANX1^CT^ or GST alone with a hydrophobic membrane spotted with 15 different membrane lipids. After washing off excess unbound PANX1^CT^, we probed the membrane using an anti-GST (Figure 3B) or anti-Panx1^CT^ antibody (Figure 3C). We did not detect an interaction between PANX1^CT^ and cholesterol; however, we found that PANX1^CT^ interacted with PA, PI(4)P, and PI(4,5)P_2_.

**Figure 3.**
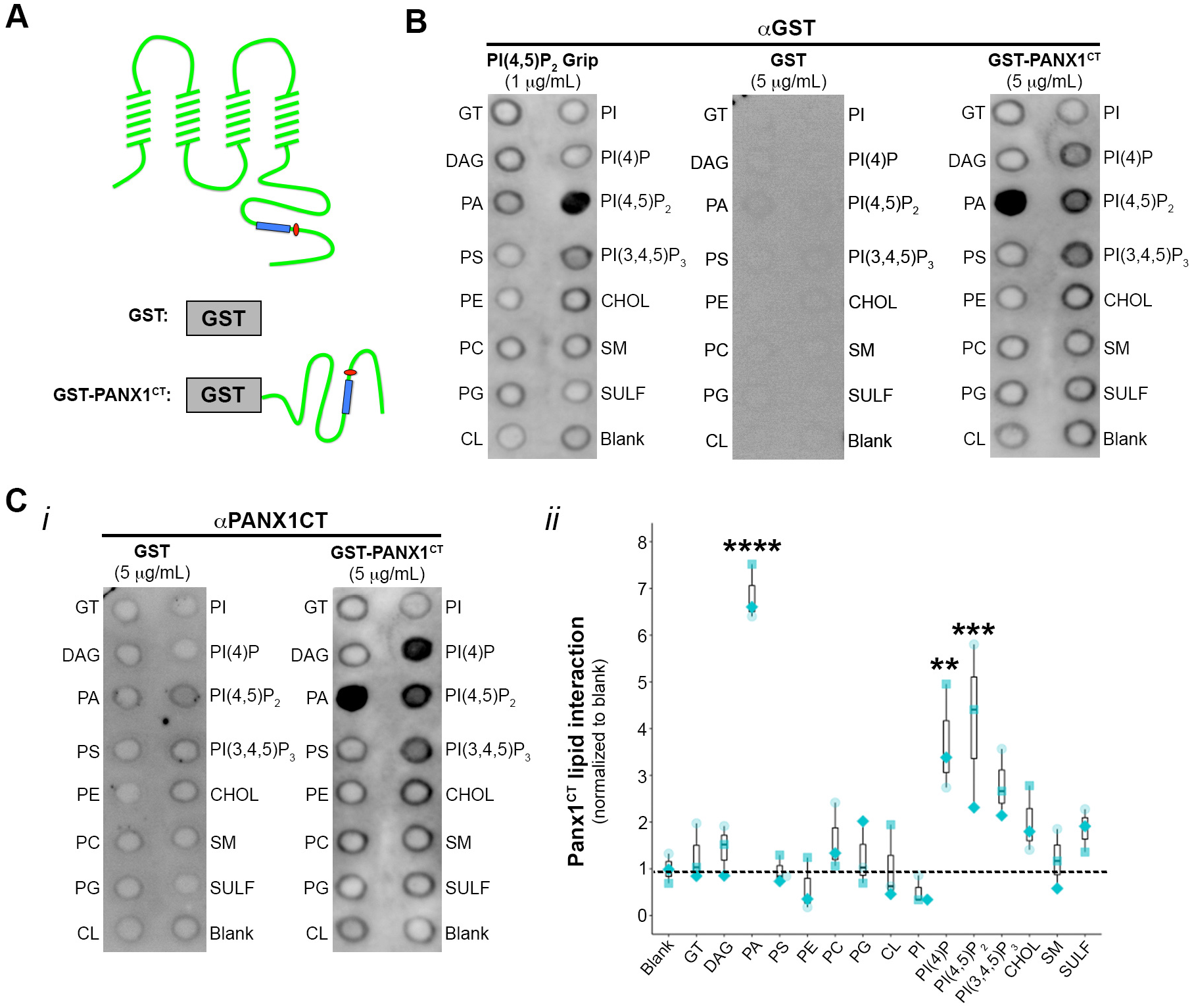
The Panx1 C-terminus interacts with phospholipids involved in endocytosis. (**a**) Schematic of (***i***) full-length Panx1 as well as (***ii***) GST and GST-fused Panx1 C-terminus (GST-Panx1^CT^) indicating location of putative lipid interaction domain (blue rectangle) relative to the caspase cleavage site (red circle). (**b**) Membrane Lipid Interaction Strip (hydrophobic membrane spotted with 15 different membrane lipids [GT – glyceryl tripalmitate, DAG – diacylglycerol, PA – phosphatidic acid, PS – phosphatidylserine, PE – phosphatidylethanolamine, PC – phosphatidylcholine, PG – phosphatidylglycerol, CL – cardiolipin, PI – phosphatidylinositol, PI(3)P – phosphatidylinositol 3-phosphate, PI(4,5)P_2_ – phosphatidylinositol 3,4-bisphosphate, PI(3,4,5)P_3_ – phosphatidylinositol 3,4,5-triphosphate, CHOL – cholesterol, SM – sphingomyelin, SULF – 3-sulfogalactosylceramide] and a blank [xylene cyanol FF]) incubated with purified peptides including positive control (PI(4,5)P2 Grip (1 μg/mL)), GST (5 μg/mL) or GST-Panx1^CT^ (5 μg/mL) and probed with GST antibody. (**c**) Membrane Lipid Interaction Strip (***i***) that was pre-incubated with purified GST (5 μg/mL) or GST-Panx1^CT^ (5 μg/mL) and probed with Panx1^CT^ antibody. (ii) Relative GST-Panx1^CT^ lipid interaction enrichment on membrane lipid interaction strip, normalized to blank, was quantified using densitometry. *N* = 3 coverslips from independent cultures (large symbols), technical replicates per coverslip are displayed in distinct shades of the same colour as smaller symbols, one-way ANOVA with Dunnett’s *post-hoc* (**p<0.01, ***p<0.001, ****p<0.0001). This figure was modified from the PhD thesis of A.K.J.B [67].

### Cell area expansion triggered by extracellular ATP is prevented by amiloride and PANX1W74A expression

Growth factor-stimulated macropinocytosis is known to increase cell size (e.g. nerve growth factor [76], insulin-like growth factor [77], for review see Lloyd et al. [64]). We found that cross-sectional cellular area was increased in cells expressing constitutively active ARF6 Q67L, independent of ATP treatment, suggesting that macropinocytosis increases N2a cell area (Figure 4A). Elevated extracellular ATP expanded cross-sectional cell area (Figure 4A), while no change was observed in response to elevated extracellular ATP with pre-treatment of amiloride (Figure 4B) or with PANX1 W74A expression (Figure 4C), suggesting that PANX1 itself might regulate macropinocytosis.

**Figure 4.**
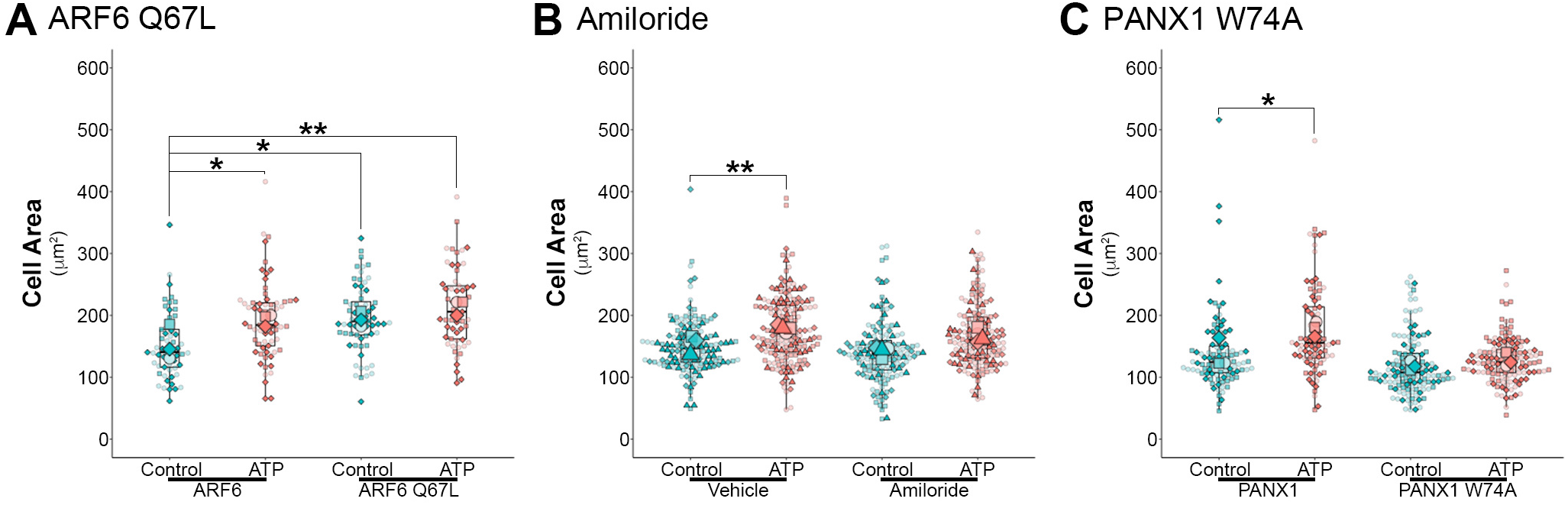
ATP-dependent macropinocytosis drives an increase in cell size that is disrupted by the W74A mutation in PANX1. (**a**) Quantification of cross-sectional cellular area in PANX1-RFP N2a cells co-expressed with ARF6-CFP or ARF6 Q67L-CFP following vehicle (water) or ATP (500 μM; 30 min; normalized to vehicle treated ARF6-CFP). Data were subjected to one-way ANOVA with Dunnett’s *post hoc* (*N* = 3; *p<0.05, **p<0.01) (**b**) Quantification of cross-sectional cellular area in PANX1-EGFP N2a cells pre-treated with vehicle (DMSO) or amiloride (300 μM) prior to vehicle (water) or ATP (500 μM; 30 min). Data were subjected to one-way ANOVA with Dunnett’s *post hoc* (*N* = 4; **p<0.01). (**c**) Quantification of cross-sectional cellular area in PANX1-EGFP or PANX1 W74A-EGFP N2a cells following treatment with vehicle (water) or ATP (500 μM). Data were subjected to one-way ANOVA with Dunnett’s *post hoc* (*N* = 4; *p<0.05). Biological replicates (*N*) are coverslips from independent cultures (large symbols), technical replicates per coverslip are displayed in distinct shades of the same colour as smaller symbols.

## DISCUSSION

Our data presented here suggest that ATP-induced PANX1 internalization occurs through macropinocytosis. Using constitutively active ARF6 Q67L we confirmed that macropinocytosis causes an expansion of N2a cell area. Similarly, ATP triggered an increase in cell area that was prevented by amiloride pre-treatment, suggesting that ATP induces macropinocytosis in N2a cells, as has been reported in other cell types. Notably, PANX1 W74A, which disrupts the ATP-dependent interaction with P2X7R and downstream internalization [14,60], also eliminated the increase in cell area triggered by extracellular ATP. Recently published structures of PANX1 at atomic resolution demonstrated that W74 and its neighbouring residue R75 form the anionic selectivity filter for PANX1 [38–40,42], likely through the formation of a cation-π interaction that stabilizes both large and small anionic molecules as they move through the narrowest portion of the pore [42]. Alanine substitution of this residue leads to structural change in the extracellular domain and loss of the cation-π interaction and loss of anion-specific selectivity [38,42], allowing for cation permeation [38,42], and loss of carbenoxolone sensitivity [38]. We and other have also demonstrated that mutation of this residue disrupts ATP-mediated PANX1 inhibition [57–59] and P2X7R-interaction dependent internalization [14,60]. Whether PANX1 W74A’s disruption of ATP-induced cell area expansion results from impaired PANX1 internalization, disrupted macropinocytosis, or changes in its channel properties is not yet clear.

Precise regulation of purinergic signalling via PANX1 is critical in many diverse cell types, including maintaining neural stem cell populations [78–80], regulating the development of immature neurons [81,82], tumorigenicity [83,84] and metastasis [85] of several cancers [86]. Macropinocytosis is also intimately involved in regulating these processes ([3–11], reviewed in Finicle et al. [12] and Bloomfield and Kay [13]). ATP-mediated macropinocytosis might not only change the receptor contribution but could also be exploited as an adaptive response for uptake of ATP and other nutrients in cells with elevated energy requirements (*i.e*., cancer [10,12]). Macropinocytosis activity is elevated at the core of tumours where nutrients were least concentrated [87]. In several types of human lung cancer (as well as breast, liver, and pancreatic cancers [88]), macropinocytosis is used as a mechanism of uptake for ATP, seen by colocalization of fluorescently tagged ATP and 70 kDa dextran [88–92]. Here, internalized ATP is used as an energy source and promotes metastasis in nutrient-starved cancer cells [92]. Moreover, constitutive macropinocytosis promotes cell proliferation in many forms of cancer [93] (reviewed in Stow et al. [94]).

ARF6 is a critical macropinocytosis effector [69,70] and a driver of macropinosome trafficking following internalization [17,18,74]. Not only did the constitutively active GTP-hydrolysis resistant ARF6 Q67L mutant drive an increase in intracellular PANX1 in macropinosomes under basal conditions (Figure 2), it also caused an expansion of cell size (Figure 4). Elevated ATP did not cause a further increase in intracellular PANX1 or cell size during ARF6 Q67L suggesting that membrane internalization was saturated under these conditions. PANX1 recycling in the presence of ARF6 and its corresponding mutants was not assessed during these experiments and will be the focus of future studies. In further support of an ARF6-dependent mechanism, PI(4,5)P_2_ and PA, PANX1^CT^ lipid interactors, are produced in response to ARF6 activation and critical for multiple stages of macropinocytosis [17], as described above and reviewed in Donaldson *et al.* [95]. Additional confirmation, such as liposome co-sedimentation assays [96], will be the focus of future work.

PANX1 demonstrates cell type-specific mechanosensitivity [44,49,52,97], where changes in cell volume or mechanical stress activate PANX1 channels. In oocytes and HEK293T cells, osmotic shrinkage activates PANX1-mediated membrane currents (but no dye uptake) while swelling does not impact PANX1 function [44]. While in airway epithelia, hypotonic challenge stimulated ATP release and dye-uptake [52]. PANX1 activation from cell shrinkage [44] suggests that ATP release by PANX1 may act in a homeostatic autocrine manner to normalize cell size. Not only does PANX1 impact cell size, there is also potential that, once internalized, PANX1 could regulate endosomal volume. Channel-dependent efflux of Cl-from endosome compartments was recently demonstrated to regulate endosome volume [98], thus the anion-selective PANX1 may be able to regulate endosome volume, if active, following internalization. Future investigations of the functional consequences of the relationship between PANX1 mechanosensation and macropinocytosis, and the function of endosome resident PANX1 could resolve these outstanding questions.

Taken together, our data suggested that ATP-dependent PANX1 internalization involved macropinocytic machinery and that PANX1 also regulates macropinocytosis, itself. The coordinated relationship between PANX1 and macropinocytosis has implications for many cellular behaviours where purinergic signalling is involved, particularly those, like cancer, where ATP can act as both a signalling molecule/growth factor and as an energy source, if internalized.

## ACKNOWLEDGEMENTS

This work was supported by operating grants from the Natural Sciences and Engineering Research Council of Canada (RGPIN-2017-03889), from the Canadian Institutes of Health Research (MOP142215), and the University of Victoria-Division of Medical Sciences to L.A.S. L.A.S. was also supported by a Michael Smith Foundation for Health Research and British Columbia Schizophrenia Society Foundation Scholar Award (5900). A.K.J.B. was supported by scholarships from NSERC (PGSD 459931-2014) and the University of Victoria (President’s Research Scholarship, Dr. Howard E. Petch and Dr. Julius F. Schleicher Memorial Scholarships) for this work; and was recently supported by the Donald Burns & Louise Berlin Postdoctoral Fellowship in Dementia Research. J.C.S.A. was supported by a University of Victoria Fellowship Graduate Award. L.A.S. is also grateful for infrastructure support from the Canada Foundation for Innovation (29462) and the BC Knowledge Development Fund (804754) for the Leica SP8 microscope system. These data are included in the PhD thesis of A.K.J.B [67].

## AUTHOR CONTRIBUTIONS

A.K.J.B. and L.A.S. conceived of the studies. A.K.J.B. performed the experiments. A.K.J.B. and E.V.D.S. performed the analysis with input from J.C.S.A and L.A.S. A.K.J.B. wrote the first draft and A.K.J.B., L.A.S., E.V.D.S. and J.C.S.A. revised the manuscript.

